# The *Mycobacterium tuberculosis* transposon sequencing database (MtbTnDB): a large-scale guide to genetic conditional essentiality

**DOI:** 10.1101/2021.03.05.434127

**Authors:** Adrian Jinich, Anisha Zaveri, Michael A. DeJesus, Emanuel Flores-Bautista, Ricardo Almada-Monter, Clare M. Smith, Christopher M. Sassetti, Jeremy M. Rock, Sabine Ehrt, Dirk Schnappinger, Thomas R. Ioerger, Kyu Y. Rhee

## Abstract

Characterizing genetic essentiality across various conditions is fundamental for understanding gene function. Transposon sequencing (TnSeq) is a powerful technique to generate genome-wide essentiality profiles in bacteria and has been extensively applied to *Mycobacterium tuberculosis* (Mtb). Dozens of TnSeq screens have yielded valuable insights into the biology of Mtb in vitro, inside macrophages, and in model host organisms. Despite their value, these Mtb TnSeq profiles have not been standardized or collated into a single, easily searchable database. This results in significant challenges when attempting to query and compare these resources, limiting our ability to obtain a comprehensive and consistent understanding of genetic conditional essentiality in Mtb. We address this problem by building a central repository of publicly available Mtb TnSeq screens, the Mtb transposon sequencing database (MtbTnDB). The MtbTnDB is a living resource that encompasses to date ≈150 standardized TnSeq screens, enabling open access to data, visualizations, and functional predictions through an interactive web app (www.mtbtndb.app). We conduct several statistical analyses on the complete database, such as demonstrating that (i) genes in the same genomic neighborhood have similar TnSeq profiles, and (ii) clusters of genes with similar TnSeq profiles are enriched for genes from similar functional categories. We further analyze the performance of machine learning models trained on TnSeq profiles to predict functional annotation of orphan genes in Mtb. By facilitating the comparison of TnSeq screens across conditions, the MtbTnDB will accelerate the exploration of conditional genetic essentiality, provide insights into the functional organization of Mtb genes, and help predict gene function in this important human pathogen.

## Introduction

Assessing the essentiality profile of a gene across different conditions is often a useful first step towards determining its function. When determined for all genes in an organism, similar patterns of essentiality across a set of conditions can indicate a shared or common function, as in the case of enzymes in a metabolic pathway [1]. Evidence of such shared or related functions can be further informed by constructing gene-gene interaction networks consisting of genes that are essential on background mutant strains that harbor single or multiple gene deletions [2,3]. In the context of pathogenic bacteria, genes that are essential for virulence in clinically relevant conditions represent potential high priority targets for drug development [4,5].

For many bacterial organisms, including the human pathogen *Mycobacterium tuberculosis* (*Mtb*), one of the main approaches to investigate the effect of gene knockouts on fitness at genome scale is to generate pooled transposon insertion mutant libraries and measure the relative growth or survival via transposon sequencing (TnSeq) [3,6,7]. In TnSeq, a saturated transposon insertion library is generated and cultured in a defined experimental condition. Amplification and sequencing of the output and control libraries yields, for each genomic region, a ratio of experiment-to-control library insertion counts. These insertion ratios are analyzed statistically in order to categorize genes as conditionally essential or non-essential [2,8,9].

Two decades ago, Rubin et al. [10] adapted transposon mutagenesis, coupled to microarray-based genome-wide fitness measurements, to *Mtb*. This method was subsequently modified using next-generation sequencing to allow single base-pair mapping of insertions [1,11]. Since then, TnSeq has been used to identify *Mtb* conditionally essential genes using *in vitro* cultures, *in vivo* mouse models as well as macrophage infection assays, and in a wide variety of conditions, such as different carbon sources, stress conditions, and a diverse array of *Mtb* clinical and mutant strains (Table 1) (Griffin et al. 2011; Zhang et al. 2012; DeJesus and Ioerger 2013; Zhang et al. 2013; Kieser et al. 2015; Mendum et al. 2015; Nambi et al. 2015; Korte et al. 2016; DeJesus et al. 2017; Xu et al. 2017; Mishra et al. 2017; Carey et al. 2018; Rittershaus et al. 2018; Bellerose et al. 2019; Minato et al. 2019; Smith et al. 2022; Meade et al. 2023). Unfortunately, these valuable genetic datasets remain scattered in supplementary tables throughout literature, making rapid analysis of the conditional essentiality profiles of genes of interest time consuming. Additionally, a lack of analytical standardization makes comparison difficult even when studies are identified, as it is often not clear how to properly compare essentiality calls that are made with different statistical approaches.

**Table 1:** The space of experimental conditions covered by the *Mtb* TnSeq Database.

| model systems | *in vitro* (bacterial culture) |  | *in vivo* (mouse) |  | in macrophage or DC |  |  |
| --- | --- | --- | --- | --- | --- | --- | --- |
| carbon sources | glycerol |  | cholesterol | trehalose | minimal media |  |  |
| stress conditions | antibiotics | hypoxia | amino acid starvation |  | nitrosative stress |  |  |
|  |  | low pH | low iron |  |  |  |  |
| *M. tuberculosis* strains | H37Rv |  |  |  | 8 clinical isolates |  |  |
| mouse strains | C57BL/6 | MHCII<br>-/- | C3HeB | BALB/cJ | Nos2-/- | Collaborative<br>Cross (CC) mouse<br>panel (61) | BXD recombinant<br>inbred panel (21) |
| *M. tuberculosis* knockout<br>strains (genetic interactions) | ponA1<br>ldtB<br>ponA2<br>ΔRv0307c<br>ΔRv0950c<br>ΔRv0954 |  | ΔRv1096<br>ΔRv1432<br>ΔRv1565c<br>ΔRv2680<br>ΔRv3005c<br>ctpC | otsA<br>ΔRv3594<br>ΔRv3684<br>ΔRv3717<br>ΔRv3811<br>ΔRv3916c |  | marP<br>eccD1<br>espl<br>PE35<br>PPE68 |  |

To address this gap, we compiled a wide array of publicly available *M. tuberculosis* TnSeq datasets into a single standardized, open-access database, which we call the *Mtb* transposon sequencing database (MtbTnDB) (Fig 1). The MtbTnDB contains 158 unique screens from 21 different publications and from the Functionalizing Lists of Unknown TB Entities (FLUTE) database (orca2.tamu.edu/U19), covering a range of experimental and physiological conditions (Table 1). To help facilitate access to the database for the TB research community, we developed an online tool that allows querying the database by either TnSeq screen or gene of interest and provides informative analyses and visualizations of the compiled data. We illustrate the potential value of the MtbTnDB for TB research through example applications, such as easily identifying orphan (i.e unannotated) genes that are conditionally essential in a given TnSeq screen. Finally, we apply unsupervised and supervised machine learning techniques to extract insights into the function of genes from their TnSeq profiles.

**Fig. 1:**
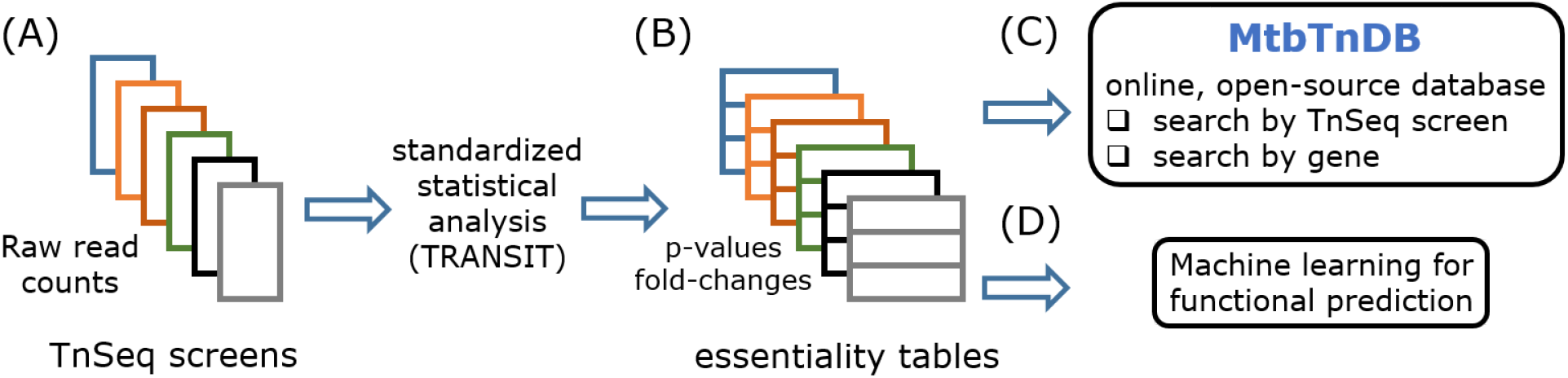
Schematic diagram of compilation and standardization of *M. tuberculosis* TnSeq screens. (A) Raw sequencing reads were collected from publicly available TnSeq screens in the literature and in the FLUTE database. (B) Using a standardized statistical processing framework with TRANSIT [9], each set of raw reads was converted into an essentiality table, with genes in the Mtb genome assigned log2 fold-change values (relative to specified control conditions) and p-values corrected for multiple hypothesis testing. (C) The set of TnSeq data can be queried in the MtbTnDB online portal, either by screen or by gene of interest. (D) The database is amenable to both supervised and unsupervised machine learning approaches to generate hypotheses about gene function.

## Material & Methods

### Standardization of TnSeq datasets

To analytically standardize and process available TnSeq raw sequencing data, we selected TRANSIT, a Python-based TnSeq analysis tool, and TPP, a software tool for processing raw reads [9]. We obtained either original raw sequencing data or read-counts in a format suitable for analysis, such as Wiggle Track Format (.wig), for 146 out of the 158 TnSeq screens in the database. For conditions where raw reads were available, we first used Tn-Seq Pre-Processor (TPP), a preprocessing tool included with TRANSIT, to map them all to the genome sequence of the standard H37Rv reference strain and obtain .wig formatted files.

Those datasets for which we could not obtain raw sequencing data (12 screens) were not included for analysis but are referenced in the MtbTnDB website to allow users to look at their results. As an additional quality control step, we only included screens for which the transposon insertion density was higher than 25%. Raw reads were mapped with TPP to the H37Rv genome using default settings. We note that for screens that did not use H37Rv, genes that are deleted in the experimental strain but not in the control strain - or vice versa - may cause false positive conditional essentiality calls. The resulting .wig files produced by TPP were analyzed using the resampling in TRANSIT, which compares two experimental conditions to find significantly different read-counts in a gene. We also included pseudocounts in the resampling pipeline, using the default of PC=1 in TRANSIT. Whenever possible, a condition (experiment) was compared to the reference dataset (control) specified in a TnSeq screen’s source publication. For publications that did not specify a reference or control condition, a highly-saturated in-vitro dataset [24] was used. Resampling was run using default settings, using TTR normalization and 10,000 permutations. P-values were automatically adjusted for multiple hypothesis testing by TRANSIT using the Benjamini-Hochberg procedure.

We labeled genes as conditionally essential in a given TnSeq screen the following two conditions are satisfied: p-val (adjusted) <= 0.05 and *abs*(log2FC)>=1, where 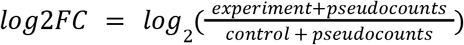, i.e. the logarithm base 2 of the ratio of insertion counts (adding the fixed so-called pseudocounts parameter), and *abs* is the absolute value. We note that the use of the absolute value in the threshold condition results in some selected genes having positive log2FC values, meaning they are detected as significantly *less* essential than expected by chance in that particular condition, with their loss providing a fitness *advantage*.

To inspect the TnSeq screens for potential batch effects, each TnSeq screen was represented by an N-dimensional vector of log2FC (where N is the number of genes in the Mtb genome) and we performed dimensionality reduction of these vectors using the uniform manifold approximation and projection (UMAP) algorithm.

### MtbTnDB website and open-source code

The MtbTnDB online portal was built using Dash (version 1.14.0), a tool for web-based analytical apps in Python. All source code used to build the online portal as well as all analyses mentioned in this work are available at https://github.com/ajinich/mtb_tn_db_demo.

### Classification of genes into annotation categories

We classified genes in the H37Rv genome into a set of five categories ranging from least to most well characterized using annotation scores from the UniProt Knowledgebase (UniProtKB) [25], which evaluates available experimental evidence for each gene at the protein level.

### Clustering of genes with UMAP

To perform the unsupervised learning analysis of the standardized TnSeq profiles using UMAP, we first removed all genes (2453) that have zero conditional essentiality calls across all 146 standardized screens. We then used the standardized binary TnSeq profiles for the remaining number of genes input into UMAP (with 3 components), as implemented in Python. Having obtained the UMAP components, we performed K-means clustering in the 2-dimensional projection, selecting the optimal number of clusters that maximizes the Silhouette Coefficient [26]. We used Fischer’s (hypergeometric) enrichment test to look for potential enrichments of COG or Tuberculist annotation functional categories within each cluster. Finally, to further evaluate the statistical significance of the observed enrichments within clusters, we generated randomized versions of the UMAP dataset by fixing the UMAP coordinates and cluster identities, shuffling the gene identifiers and annotation information associated to each point in the UMAP projection, and repeating the statistical enrichment test.

### Prediction of gene function

To fit a supervised machine learning model to the standardized TnSeq profiles, we used the log2FC values from the standardized matrix as inputs. Genes containing missing values in some screens were dropped (n=84). We used Tuberculist annotation functional categories as classification labels and dropped genes belonging to the ‘unknown’ (n=14) or ‘conserved hypothetical’ (n=1020) categories. The resulting matrix containing 2937 genes was used for prediction.

We used two sub-models to predict functional categories: a logistic regression classifier with L1 regularization and a random forest classifier [27]. Both methods were used as implemented in the scikit-learn library (version 0.23.0) in Python with default parameters except for the L1 regularization strength, C, which was set to 10. We used SMOTE [44] to correct for class imbalance along with 3-fold cross-validation using stratified splits. The outputs of each of the sub-models were subsequently fit to a logistic regression classifier (default parameters, except C=10) to create an ensemble model for final predictions. The ensemble model showed better performance than the sub-models run individually.

## Results

### Compiling and standardizing *M. tuberculosis* TnSeq datasets

We set out to build a database that compiles all publicly available *M. tuberculosis* TnSeq datasets in a standardized, open-source format (Fig. 1). We assembled data for *Mtb* TnSeq screens encompassing a total of 158 comparisons between experimental and control conditions from two main sources. A first set of 143 comparisons were compiled from 21 publications (Griffin et al. 2011; Zhang et al. 2012; DeJesus and Ioerger 2013; Zhang et al. 2013; Kieser et al. 2015; Mendum et al. 2015; Nambi et al. 2015; Korte et al. 2016; DeJesus et al. 2017; Xu et al. 2017; Mishra et al. 2017; Carey et al. 2018; Rittershaus et al. 2018; Bellerose et al. 2019; Minato et al. 2019; Sassetti et al. 2003; Sassetti and Rubin 2003; Rengarajan et al. 2005; Joshi et al. 2006; Smith et al. 2022; Meade et al. 2023). The remaining 15 were obtained from the Functionalizing Lists of Unknown TB Entities (FLUTE) database (orca2.tamu.edu/U19) .

The set of unique TnSeq screens in the MtbTnDB cover a wide range of different experimental conditions (Table 1). Of the 158 screens in the database, 47 are *in vitro* bacterial culture experiments, covering many different culture media (carbon sources and limiting nutrients) and stress conditions (antibiotics, low oxygen tensions, acidic pH, nitrosative stress, iron limitation, and amino acid starvation). A second set of 109 screens are *in vivo* experiments using mouse infection models. Two recent publications (Meade et al. 2023; Smith et al. 2022) are the source of the large majority (82 out of 109) of these *in vivo* screens. An additional 2 TnSeq screens use cell culture (macrophage and dendritic cell) infection models. Although the large majority of the experiments were performed using the standard laboratory Mtb strain (H37Rv), one publication (Carey et al. 2018) performed comparative screens on eight different clinical isolates. A number of the TnSeq comparisons (34) were performed on *Mtb* strains harboring a single gene knockout, yielding information on genetic interaction networks underlying observed conditional essentiality profiles. Finally, for completion we also included data for 5 screens from the literature that were performed using microarray-based transposon site hybridization (TraSH) [28–31].

### Standardized statistical analysis of TnSeq screens using TRANSIT2

In order to facilitate comparative analyses across different TnSeq screens, we set out to re-process the raw data for as many screens as possible using a single standardized statistical method based on TRANSIT, a Python-based TnSeq analysis tool (see Methods for details). Although TRANSIT accommodates a variety of statistical approaches for making essentiality calls, we utilized the resampling method to compare every experimental condition to an appropriate reference or control condition. Briefly, resampling compares read-counts between two different conditions utilizing a permutation test to identify significant differences in the mean read-counts [9]. Resulting hits are labeled as *conditionally* essential genes: genes that are essential in one background condition (experiment) but not another (reference or control). By comparing against a suitable reference or control condition, we sought to reduce potential biases that can exist due to differences in mutant libraries. Whenever possible, the reference condition was chosen to be the one described in the original TnSeq screen. When no suitable reference or control condition was specified or included in the screen, we utilized a fully saturated in-vitro dataset [24] as the control condition.

In order to analyze the effect of our standardized processing of the underlying TnSeq sequencing data, we compared the number of conditionally essential genes detected in each screen (using fixed threshold values for the corrected p-value<=0.05, and a log(2)-fold change>=1) against those reported in the corresponding original, individual datasets (Table S1). Across all screens analyzed, we find that the standardized data and the original analyses differ in around 2% of the conditional essentiality calls. Approximately half of these are cases where the standardized dataset suggests that a gene is conditionally essential in a particular screen, whereas the original analysis did not, and the other half of the cases correspond to the opposite situation. A few individual screens have a relatively high number of non-consensus essentiality calls between the standardized and original datasets. In particular, one TnSeq screen in an *in vivo* mouse model [13] has 403 genes that were called essential in the original datasets, but not in the standardized analysis. One possible reason behind this large inconsistency is the different statistical methods used to analyze the sequencing data. We note that both the original and the standardized data sets are included in the MtbTnDB (see section on online database).

In addition, we inspected the standardized TnSeq screens in search of batch effects, i.e. variations in the datasets that occur independently of biological signal and noise. Dimensionality reduction of data vectors representing each TnSeq screen (Methods) revealed that, of the 76 screens included in the database, a set of eight TnSeq screens formed a distinctly separate cluster (Fig. S1). This cluster consists of screens where essentiality counts in eight clinical strains are compared relative to a control strain (H37Rv), which could either be indicative of experimental batch effects or true biological differences between domesticated laboratory strains and clinical isolates.

### The statistics of conditional essentiality across the MtbTnDB

Fig. 2A shows the distribution of conditional essentiality calls across all genes in the *Mtb* genome. We find that 2453 genes are not labelled as conditionally essential in any of our standardized screens. We note that a significant subset of these (425) are genes that are essential or have a significant growth defect in the fully saturated *in-vitro* screen [24] that we used as a control (reference) condition for a large number of comparisons. Thus, despite being essential for growth in standard in-vitro culture conditions, these genes are not detected as conditionally essential in any screen in the MtbTnDB: i.e there is no significant difference in their degree of essentiality in standard in-vitro culture in comparison to the experimental conditions covered in the database. The rest of the genes with zero conditionally essential calls likely represent a mixture of putatively dispensable genes and (i) genes with essential roles in experimental conditions that are outside the coverage of our database, (ii) false negatives, i.e. genes that do not meet the threshold of statistical significance due to experimental factors like low coverage.

**Fig. 2:**
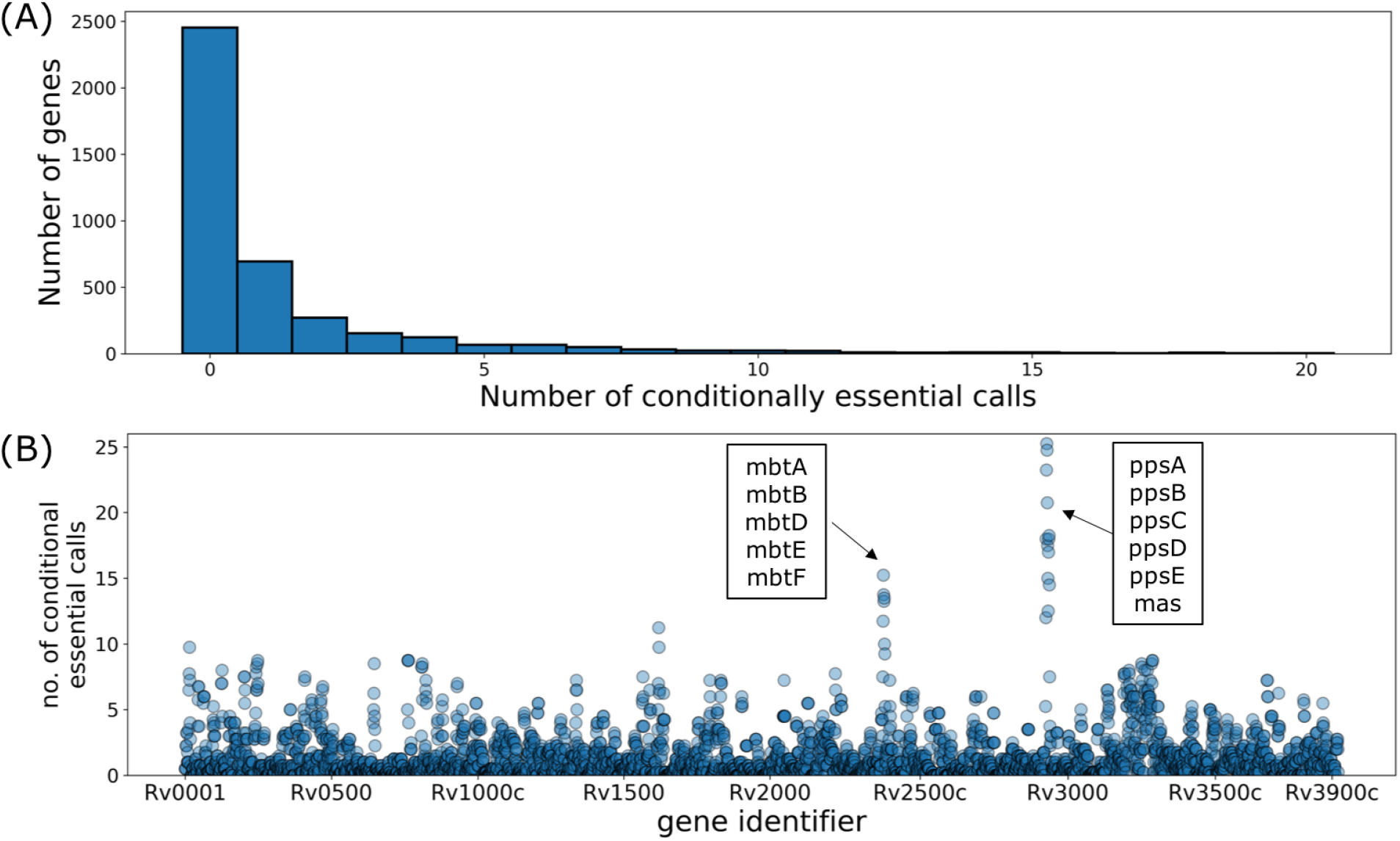
Distribution of conditional essentiality calls in the MtbTnDB across all genes. (A) Distribution of the number of conditionally essential calls across all genes in the *Mtb* genomes (B) Number of conditional essentiality calls vs. position in the genome across the MtbTnDB for all H37Rv *Mtb* genes. The mycobactin and PDIM biosynthetic operons, which tend to have a large number of conditionally essential calls, are highlighted.

At the tail of the distribution, we find approximately 120 genes that are conditionally essential in 10 or more comparative TnSeq screens. Among these (Fig. 2B) we find a strong enrichment for phthiocerol synthesis polyketide synthase type I genes (*ppsA-E*), with all members of these PDIM biosynthetic pathway genes having at least 24 conditional essentiality counts. One possible explanation for this is the fact that loss of PDIM biosynthesis genes *in vitro* confers a fitness advantage to *Mtb* [32]. The fully saturated in-vitro dataset [34] that was used as the control condition for a large number of comparisons contains PDIM biosynthesis genes, but some of these genes could have been lost in several of the experimental conditions, leading to few or no detectable transcription insertion counts and resulting in false positive conditionally essential call. A second set of genes with a high number of conditionally essential calls are the members of the mycobactin siderophore biosynthetic pathway (*mbtA, mbtB, mtbD, mbtE, mbtF*) [33] (Fig. 2B). Interestingly, in more than half of these instances the mycobactin biosynthesis genes are detected as a significant hit because they are *less* essential (i.e. p-val (adjusted) <= 0.05 and log2FC >= 1) implying their loss provides a fitness advantage across a range of different conditions. One might expect that iron-scavenging would not be required across several *in-vitro* conditions, as most media used in TnSeq experiments is replete with iron.

### MtbTnDB, an online database of standardized Mtb TnSeq screens

To allow efficient querying of the standardized data, we developed an MtbTnDB web interface (www.mtbtndb.app). The interface allows three methods of querying the database - by screen (Fig. 3), by gene, and in co-essentiality mode. Each screen is a pairwise comparison between an experimental and control condition. The “Analyze datasets’’ tab provides an information-rich and interactive overview of the selected comparison. An “About this dataset” section includes a description of the screen, links to the original publication, and the number of replicates used for both the experimental and control conditions. A volcano plot and accompanying table allow interrogation into differentially represented mutants. Sliders for significant log2FC and qval cut-offs allow interactive coloring of data points on the volcano plot. Mutants of interest can also be highlighted on the volcano plot by selection in the accompanying table. We also provide two plots depicting broader analyses of the data. First, a bar plot depicts the enrichment of COG functional categories in over/under-represented mutants. Second, a bubble plot allows identification of the subset of genes that are both conditionally essential in this analysis and the least annotated subset, providing useful pointers for follow-up experiments. An example case using this type of plot is provided in the next section.

**Fig. 3:**
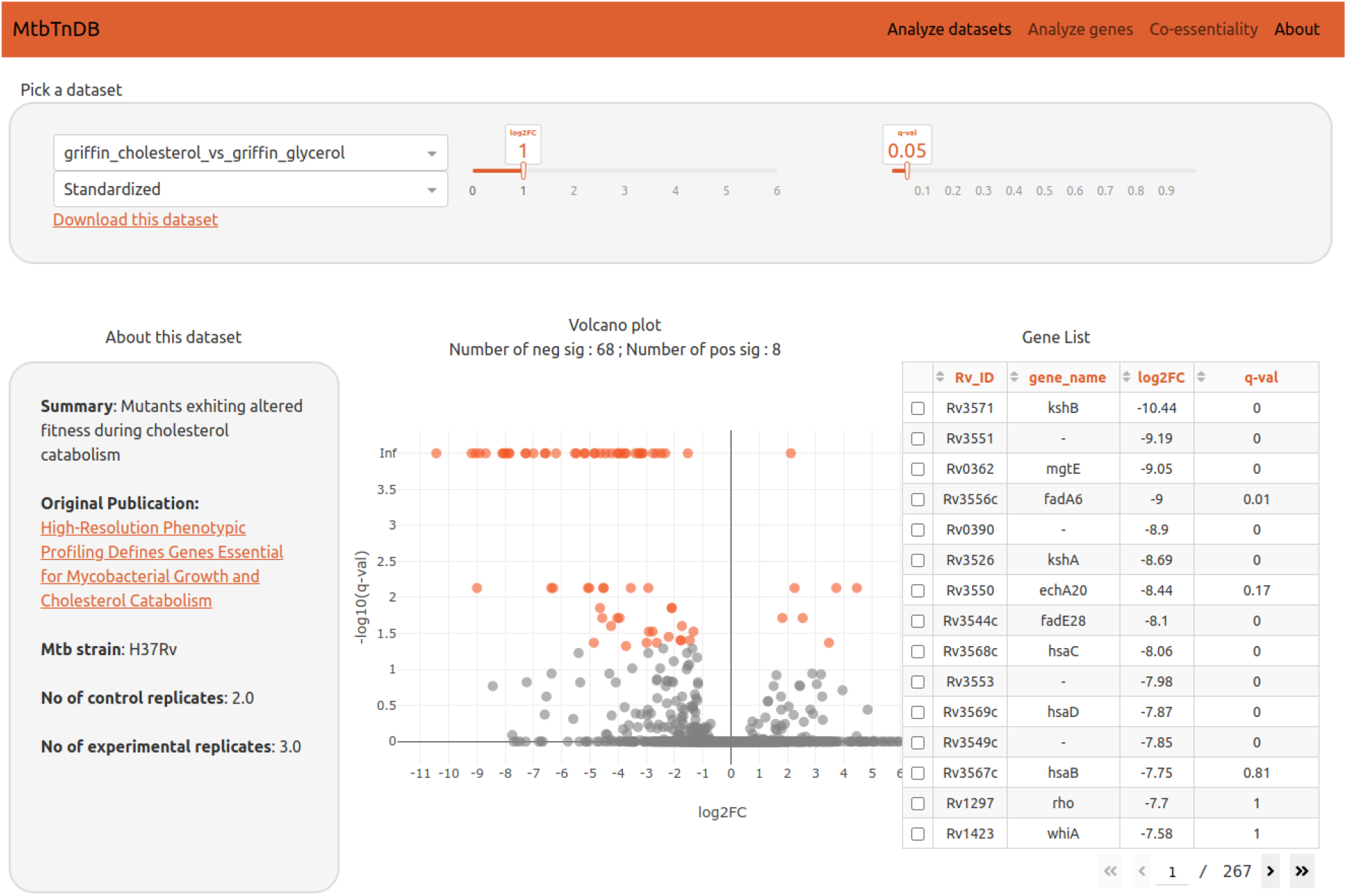
An online explorer of TnSeq datasets. A screenshot of the “Analyze datasets” modality is shown. Users can select which TnSeq screen to explore, whether to display either the standardized or the original publication’s dataset, as well as significance thresholds for log2 fold-changes and q-values. A brief summary of the TnSeq screen is shown in the left panel, alongside a link to the original publication and the number of replicates for both control and experimental conditions. An interactive volcano plot highlights conditionally essential genes, which can also be accessed and selected through the table on the right.

The second functionality provided in the web interface is to query genes of interest and list the set of TnSeq screens in which mutants for this gene are differentially represented. In addition to log2FC and q-values we also provide track-views, which are visual depictions of the magnitude and location of insertions within the selected gene.

The third tab in the MtbTnDB web interface serves as a dedicated co-essentiality analysis feature, which identifies significant correlations between the TnSeq profiles of gene pairs — specifically, the log2-fold changes — across all experimental screens within the database. This functionality allows researchers to select a gene of interest and engage with an interactive visualization that shows significant correlations with other genes, suggesting potential functional interactions and dependencies. This visual and interactive interface displays both first and second-degree co-essential relationships, providing a comprehensive map of potential genetic interactions.

The data used in MtbTnDB is provided in the “About” tab, and users can also choose to download individual screens from the “Analyse datasets” tab. We also note the modularity of the database and its ability to accommodate additional datasets in the future, potentially including conditional essentiality screens using CRISPRi.

### Example usage: selecting experimental conditions to study unannotated genes

A large fraction of genes in the *Mtb* genome remains unannotated, and the MtbTnDB could help annotate such “orphan” genes by guiding the design of experimental conditions in which to assay their function. This is motivated by the notion that experimental conditions in which a mutant strain displays a growth or survival defect are more likely to reveal the orphan gene’s physiological role. For instance, if an enzyme of unknown function participates in cholesterol catabolism, the probability of discovering its function will be highest if the mutant strain’s metabolic profile is obtained when grown with cholesterol as its primary carbon source.

To help select experimental conditions in which to study orphan genes, we used experimental annotation scores from Uniprot, consisting of 5 categories ranging from least to most well characterized (see Methods), to classify genes. For every TnSeq screen in the MtbTnDB, we then obtain the unknown essentials: the set of genes that are both conditionally essential (in that experimental screen) and fall in the category of least well characterized genes (Fig. 4, top-left corner). The number of unknown conditional essentials varies across each TnSeq screen in the MtbTnDB, with 10 screens having more than 30 conditionally essential genes with the lowest annotation score. In particular, two mouse *in vivo* screens have more than 65 genes that fall in the most poorly annotated category, underscoring the fact that the function of many potential drug targets remains to be discovered. This also illustrates how the MtbTnDB can be used to systematically select experimental conditions that result in measurable growth phenotypes for orphan genes of interest for further study.

**Fig. 4:**
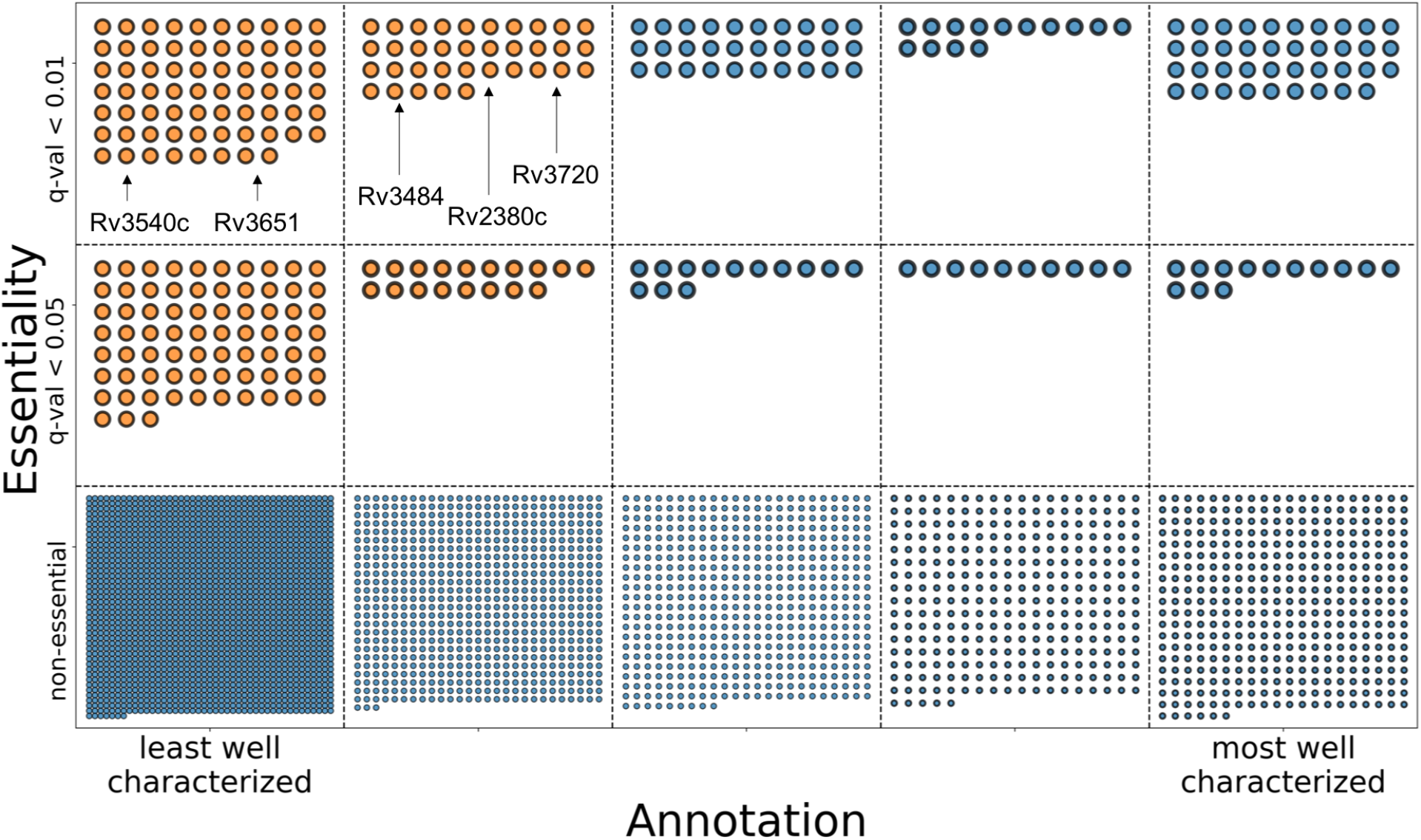
Mapping out conditionally essential genes of unknown function. Data shown corresponds to a single TnSeq screen in the MtbTnDB: an H37Rv *in vivo* (mouse) screen [2]). Genes are represented as circles, and are binned along the x-axis into five distinct categories according to their annotation level and along the y-axis according to their conditional essentiality (the exact position within each bin/category is not meaningful). The least well characterized genes (i.e orphans) that are conditionally essential in this particular TnSeq screen are shown in orange (the gene identifiers for five example genes are shown).

### Predicting functional categories with the MtbTnDB

We asked whether similar TnSeq conditional essentiality profiles are indicative of similar biological function, where a conditional essentiality profile is defined as the set of (conditional) essentiality calls across all standardized TnSeq screens; i.e. a 76-dimensional vector for each gene in the *Mtb* genome. To explore this, we applied a uniform manifold approximation and projection (UMAP), a non-linear dimensional reduction tool [34], to the standardized TnSeq data. We hypothesized that, since genes belonging to the same operons are functionally related, neighboring genes in the genome are likely to have more similar TnSeq conditional essentiality profiles than random pairs of genes on average (Fig. 5A). To test this, we compared the distribution of Euclidean distances in the UMAP projection between (i) pairs of genes that are in the same genomic neighborhood (i.e. apart from each other in the genome by a fixed cutoff genomic distance) and (ii) pairs of genes chosen at random from the full *Mtb* genome. We indeed find that neighboring genes tend to have more similar TnSeq profiles than expected by chance (p=10^-14^, Kolmogorov-Smirnov test).

**Fig. 5:**
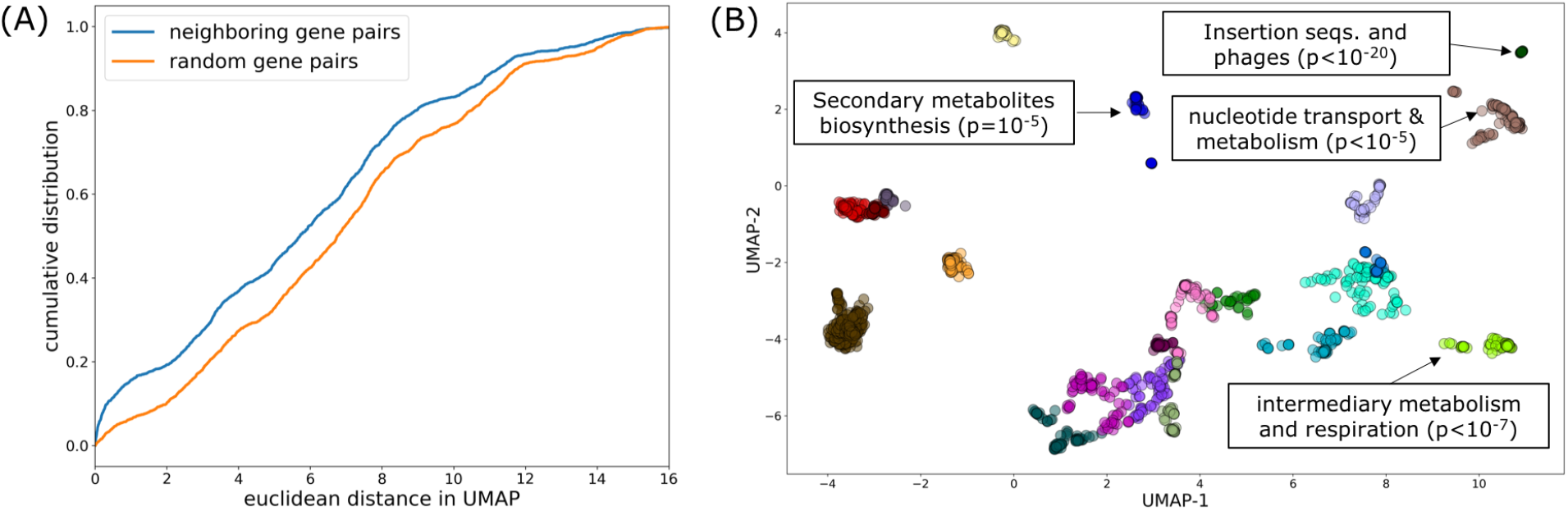
Genes with similar TnSeq profiles tend to have similar functions. Neighboring genes in the genome have more similar TnSeq profiles, and clusters of genes with similar TnSeq profiles are enriched for annotated genes belonging to the same functional categories. (A) Distribution of Euclidean distances (in UMAP projection) for pairs of random genes (orange) and neighboring genes (i.e. less than 3 genes away from each other in the genome). The two distributions are significantly different (p<10^-6^, Kolmogorov-Smirnov test). (B) Uniform manifold approximation and projection (UMAP) of TnSeq profiles. Genes with at least one conditional essentiality call in the MtbTnDB were clustered using k-means according to their UMAP coordinates and color-coded. Four clusters enriched for gene functional categories are shown.

Next, using k-means clustering, we grouped genes by the similarity of their TnSeq profiles as captured by their coordinates in the UMAP projection. An example UMAP projection with 19 different clusters is shown in (Fig. 5B). Using the COG [35] and Tuberculist [36] *Mtb* genome annotations, we find that several clusters are significantly enriched for a number of different functional categories, including, “nucleotide transport and metabolism”, “intermediary metabolism and respiration”, and “secondary metabolites and biosynthesis”, among others. Importantly, our observation of clusters enriched for functional categories is statistically significant (p<0.05, permutation test), and on average, zero categories were enriched when annotations were shuffled and re-analyzed for overlap with the UMAP clusters (Fig. S2). Taken together, these analyses serve as evidence that the TnSeq profiles capture functional relationships between genes, and encode information about the underlying operonic structure of the *Mtb* genome.

Building on the unsupervised analysis described above, we reasoned that since genes with similar physiological functions have similar essentiality profiles, the information encoded in the Mtb TnSeq Matrix could be used to help computationally annotate orphan genes. Towards this end, we trained a supervised classification model which takes as input a gene’s log2FC values from the 76 screens in the database and predicts the functional category from the Tuberculist genome annotation that the gene belongs to. Our model (see Methods), a stacked meta-learner that combines predictions from logistic regression and random forest sub-models, achieved an accuracy of 45-60% (3-fold cross-validation) for three Tuberculist categories: ‘PE/PPE’, ‘information pathways’, and ‘insertion sequences and phages’, as assessed by cross-validation (Fig. 6). We note that (i) our prediction is completely blind to the coding sequence, and that (ii) as a comparative baseline, the accuracy of an unbiased random classifier would be 12.5% for each of the 8 categories. Thus, while improvements in predictive accuracy can be achieved by incorporating other sources of data, TnSeq essentiality profiles can potentially help guide the functional annotation of unknown genes in Mtb. In addition, the predictive power of machine learning models trained on TnSeq data will likely improve as the database is populated with more conditional essentiality screens.

**Fig. 6:**
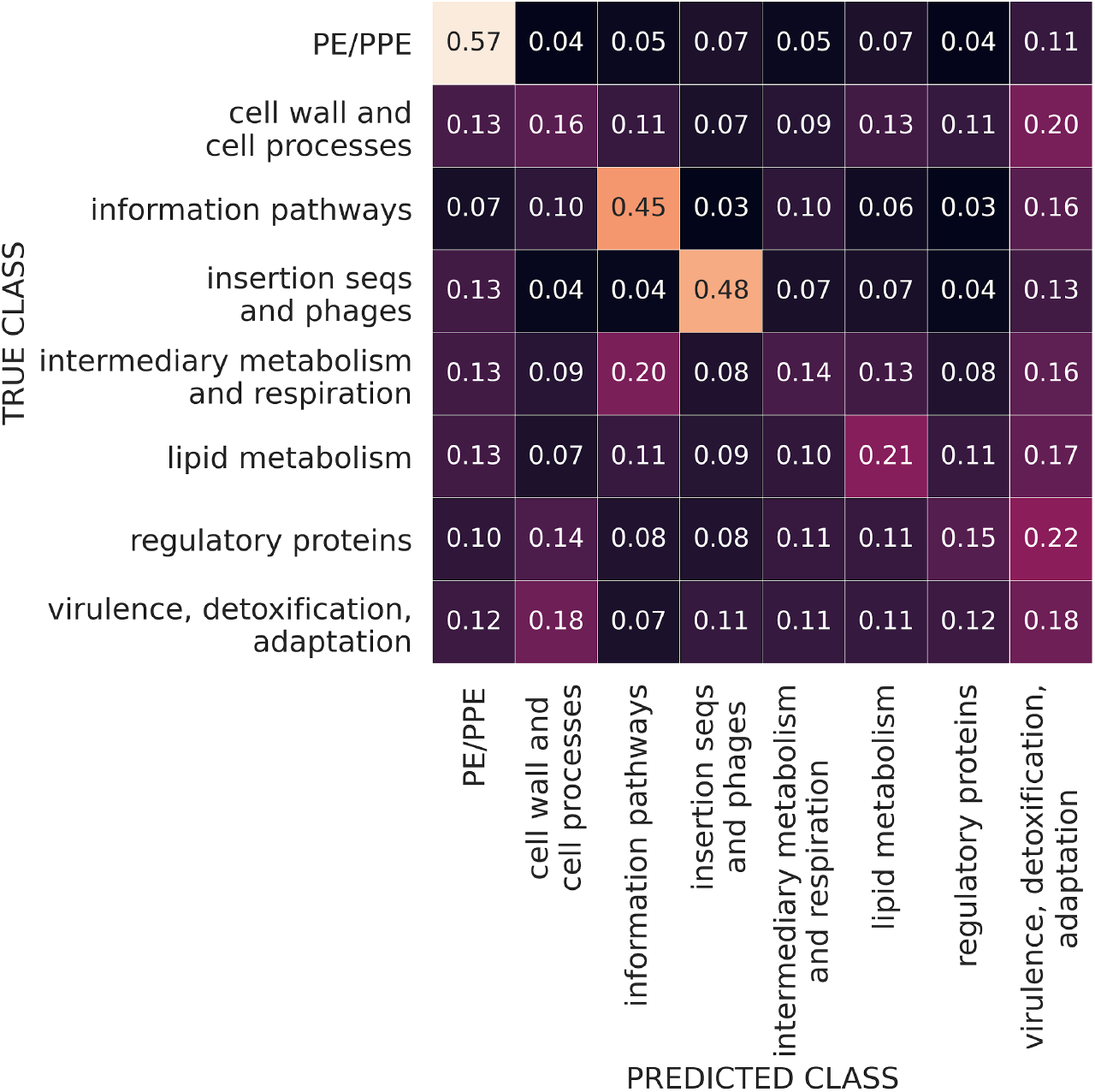
Prediction of gene function with machine learning. Confusion matrix showing the fraction of observations predicted to be classified in each functional category. Predictions were generated using an ensemble machine learning model consisting of a logistic regression and random forest classifier. The number of genes in each functional category are: PE/PPE (166); cell wall and cell processes (771); information pathways (242); insertions sequences and phages (146); intermediary metabolism and respiration (934); lipid metabolism (271); regulatory proteins (198); virulence, detoxification, adaptation (220).

## Discussion

To facilitate access to and the ability to gain insight from the vast amount of information stored in *M. tuberculosis* genetic essentiality screens, we compiled a large set of publicly available TnSeq screens into a single, standardized, and open-access database, the *Mtb* transposon sequencing database (MtbTnDB). While other TnSeq databases exist [37], these contain significantly fewer *Mtb* specific screens and do not address the issue of standardization of raw sequencing data. We envision periodically updating the MtbTnDB by adding publicly available TnSeq screens as they become available in the literature: researchers wishing to contribute their screen to the database would simply need to provide the sequence files along with a basic description of the screening conditions.

Although we aimed to maximize the degree of standardization, we note that there are upstream experimental factors that introduce variability into the screens, placing inherent limits in our ability to standardize the data. For instance, the transposon libraries differ in saturation (the number of unique insertion sites with one or more mutants present in the library), or in protocol details, such as whether barcoding was utilized, or the specific sequencing technology used to obtain the reads. These differences may introduce artifacts, such as PCR over-amplification, or simply reduce the amount of data that is available (in the case of libraries with low coverage) which ultimately will affect the statistical power of downstream analyses.

One limitation of the current set of TnSeq screens included in the MtbTnDB is a bias towards a subspace of experimental conditions. Future efforts should therefore be directed towards covering a wider and diverse region of experimental condition space. Towards this end, a recent TnSeq screen assaying iron uptake in *M. tuberculosis* [38], as well as recent technical developments that couple TnSeq with random DNA barcoding (RB-TnSeq) allowing parallel screening across dozens of conditions [39] are very encouraging.

It is worthwhile to point out that the information afforded by TnSeq experiments is quite different and complementary from that of RNAseq (transcriptomics). Transcriptomics identifies sets of genes whose expression changes in response to changes in conditions, which can be clustered into sets of co-regulated genes (or regulons) and sometimes correlated with transcription factors [2,40]. There is a large amount of transcriptomic data (DNA microarrays and RNAseq) that has been collected for Mtb in different conditions over the years, including stress conditions like hypoxia and antibiotic exposure, in macrophages and mouse models, knockout strains, among others [41]. TnSeq identifies the essentiality of genes in a given condition. In the extreme, essential genes are those which are indispensable for growth in the condition and hence cannot tolerate transposon insertions. Additionally, there might be some genes that exhibit a quantitative reduction in transposon insertion counts, which is interpreted to indicate a fitness defect (growth impairment) caused by the disruption of the ORF [28]. Hence, TnSeq reveals information about which genes play functional roles in pathways required for survival in the condition. While both technologies can be efficiently scaled up to profile genome-wide cellular responses to changes in environmental conditions, the assays assess different information about genes (essentiality versus expression), and the lists of hits (conditionally essential genes vs. differentially regulated genes) often exhibit little overlap [42]. A gene can be differentially expressed without being conditionally essential and, conversely, a gene can become conditionally essential without a significant change in expression in the new condition. Efforts to combine TnSeq and RNAseq to enhance function prediction are on-going.

The prospect of combining high-throughput phenotypic screening with machine learning to illuminate orphan gene function is promising. In this work, we tested how well the MtbTnDB could functionally classify unannotated *Mtb* genes using supervised machine learning models and found limited prediction accuracy. One possible explanation for this could be the quality of the functional annotations themselves, and future directions would include combining the MtbTnDB with complementary sources of data, such gene expression datasets, in order to improve the prediction of orphan gene function. We envision that the essentiality patterns encoded in the collected and standardized TnSeq screens will help researchers unravel the higher-order, physiological function of genes in the *Mtb* genome.

## Data availability

Software and source code is available GitHub: https://github.com/ajinich/mtb_tn_db.

## Acknowledgements

We thank Michelle Bellerose, Anthony D. Baughn, Yusuke Minato, Rainer Kalscheuer, and Kristi Guinn for assistance in providing TnSeq screen sequencing data.

