## Supplemental Figures 1 for "The *Mycobacterium tuberculosis* transposon sequencing database (MtbTnDB): a large-scale guide to genetic conditional essentiality"

### Supplementary Figures

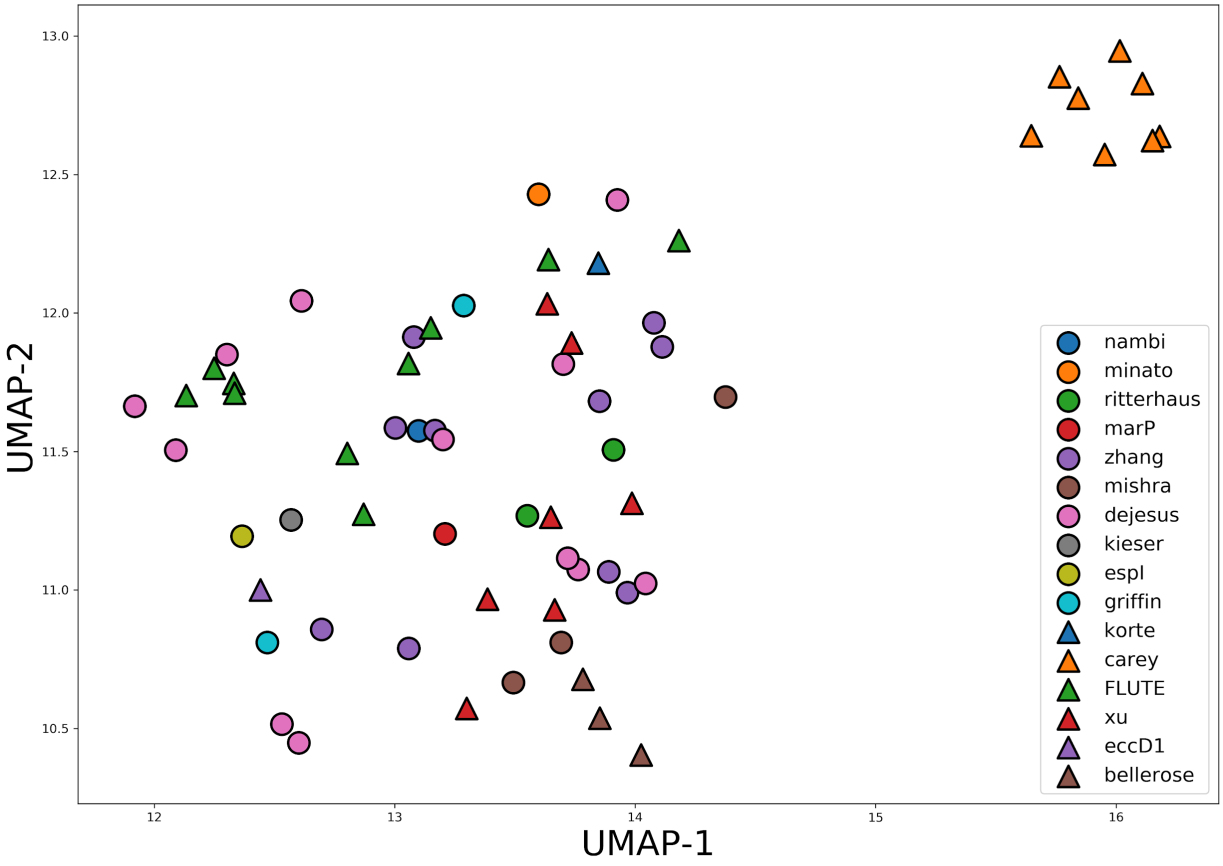


**Fig S1: Unsupervised clustering of standardized TnSeq data by screen reveals a subset of screens with potential batch effects.** Two-dimensional projection of TnSeq screens was performed using the Uniform Manifold Approximation and Projection (UMAP) algorithm. Each point corresponds to a single TnSeq screen in the MtbTnDB, and screens are color-coded according to the reference publication or data-source.

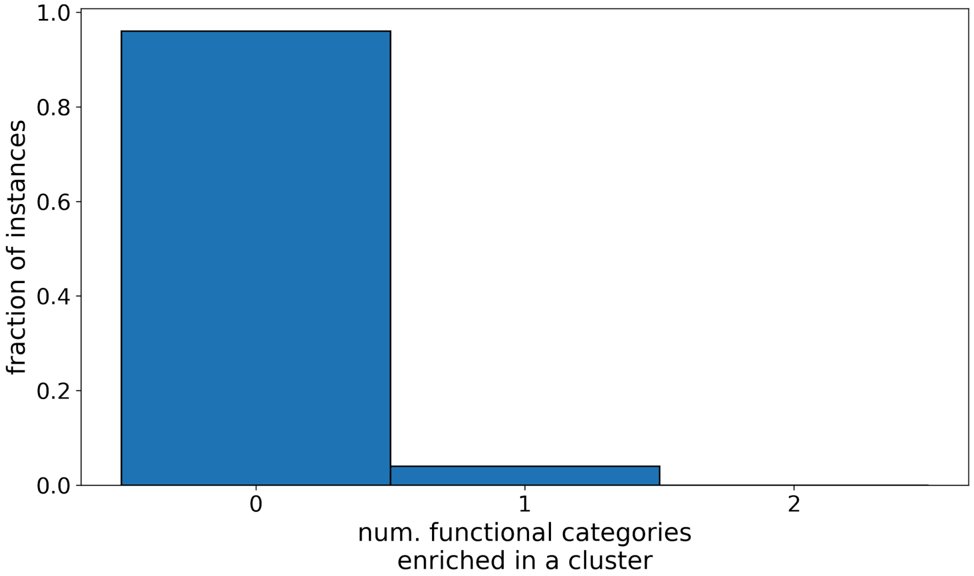


**Fig S2: Lack of enrichment of gene functional categories in randomized versions of gene essentiality data.** To evaluate the statistical significance of the observed enrichments of COG and Tuberculist gene functional categories in UMAP projection of TnSeq essentiality profiles, we generated 200 randomized versions of the UMAP dataset. Specifically, the positions of genes in the 2-dimensional UMAP projection were fixed, while the gene information (Rv-ID, name, annotation) were randomly shuffled. The normalized histogram shows that in more than 95% of the shuffled datasets, no enrichment of functional categories was detected in any cluster.

**Table S1: Consensus of conditional essentiality calls between standardized datasets and original published datasets.**

| | | standardized | |
|---|---|---|---|
| | | conditional essential | not conditional essential |
| original | conditional essential | 1157 | 1205 |
| | not conditional essential | 1292 | 107952 |
